# Kinetic control of stationary flux ratios for a wide range of biochemical processes

**DOI:** 10.1101/858126

**Authors:** Joel D. Mallory, Anatoly B. Kolomeisky, Oleg A. Igoshin

## Abstract

One of the most intriguing features of biological systems is their ability to regulate the steady-state fluxes of the underlying biochemical reactions, however the regulatory mechanisms and their physicochemical properties are not fully understood. Fundamentally, flux regulation can be explained with a chemical kinetic formalism describing the transitions between discrete states with the reaction rates defined by an underlying free-energy landscape. Which features of the energy landscape affect the flux distribution? Here we prove that the ratios of the steady-state fluxes of quasi-first-order biochemical processes are invariant to energy perturbations of the discrete states and are only affected by the energy barriers. In other words, the non-equilibrium flux distribution is under kinetic and not thermodynamic control. We illustrate the generality of this result for three biological processes. For the network describing protein folding along competing pathways, the probabilities of proceeding via these pathways are shown to be invariant to the stability of the intermediates or to the presence of additional misfolded states. For the network describing protein synthesis, the error rate and the energy expenditure per peptide bond is proven to be independent of the stability of the intermediate states. For molecular motors such as myosin-V, the ratio of forward to backward steps and the number of ATP molecules hydrolyzed per step is demonstrated to be invariant to energy perturbations of the intermediate states. These findings place important constraints on the ability of mutations and drug perturbations to affect the steady-state flux distribution for a wide class of biological processes.

## INTRODUCTION

Chemical kinetics represents a fundamental mesoscopic formalism to understand many biological phenomena with a wide range of major cellular processes described in terms of the complex networks of biochemical reactions^1^. When the mechanistic details of the process are known, the underlying mechanisms can be represented as elementary steps described by mass-action kinetics of first order reactions (e.g., for catalysis, conformational transitions, and dissociation) or second order reactions for bimolecular binding transitions. This formalism allows one to study the kinetics of enzymatically controlled reactions and resulting kinetic laws^2^, transformations between different states of promoters and resulting transcription rates^3–6^, transitions between receptor-complexes and resulting signaling^7,8^, and many other phenomena^9,10^. In many of these examples, the time-scale of changes in the concentrations of unbound molecular species is much slower than the time-scale of transitions between the states of the macromolecular systems. For example, concentrations of substrates and products of enzymatic catalysis change much slower than transitions between the different enzyme states^11^. In this regime, second-order binding reactions can be effectively approximated as quasi-first-order transitions. As a result, the kinetic equations are linear, yielding a so-called linear framework^12^. In the steady state, the resulting linear algebraic equations can be easily solved to compute probabilities, fluxes and many other important quantities.

For an arbitrary chemical network with elementary chemical transitions, the kinetics can be fundamentally understood in terms of the underlying free-energy landscape. Assuming that thermal equilibrium in each chemical state is reached much faster than the time-scale of the chemical transitions, the rate of each step can be expressed in terms of the activation energy, i.e., the difference between the energy of the state and the barrier separating the states^13^. The resulting expression of the rate constants is often referred to as the transition-state theory^14^. We note that in this picture the free energy landscape emerges naturally as a consequence of the microscopic description of a process through elementary chemical steps.

For reaction networks with cycles that do not change the concentrations of the molecular species in the cellular environment, the underlying chemical reaction rates are subject to the detailed balance constraints^15^. In that case, the resulting steady-state probabilities are computed from the Boltzmann distribution, and they only depend on the free energy of the states that are independent of the free energy barriers^15^. For example, models of binding and dissociation of transcription factors to DNA and resulting transcription rates in bacteria are often calculated in the equilibrium thermodynamics framework^16,17^. This regime is often referred to as thermodynamic control in contrast to the kinetic control regime when the barrier heights determine the resultng distributions^18^. In many other biologically relevant cases, transitions between states of the macro-molecular system are coupled with the changes in the pool of cellular cofactor molecules. If the cycles in the network change the chemical composition of the cellular environment, the system reaches a non-equilibrium steady-state where non-vanishing net fluxes are possible^15^. For example, transitions between states of enzymes and molecular motors can be coupled to the hydrolysis of ATP^19,20^. In the non-equilibrium steady state, the steady-state probabilities and the fluxes can depend on the energies of the transition states. Thus, non-equilibrium steady states may be under a combination of thermodynamic and kinetic controls, and it is not clear which features of the free energy landscape control key system properties^21^.

The non-zero values of net fluxes in the non-equilibrium steady state have important biological implications. For example, the rate of an enzymatic reaction, the speed of a molecular motor, and many other crucial characteristics are proportional to these steady-state fluxes^15,22^. Moreover, many important properties can be related to the ratios of the steady-state fluxes. Specifically, the enzyme selectivity or the error rate that quantifies the ability of enzymes to discriminate between right (cognate) and wrong (non-cognate) substrates can be expressed as the ratio of the catalytic fluxes to the corresponding states^19,21,23^. In the same way, the efficiency of a molecular motor can be expressed in terms of the ratio of forward-stepping and futile cycle fluxes^22,24^. Notably, our previous research^25^ has shown that for both Michaelis-Menten (MM) and the Hopfield kinetic proof-reading (KPR) mechanism^19^, the error rate is purely under kinetic control. As such, the error rate is only a function of the differences in the free energy barriers for pathways leading to the right and wrong substrates, and it is independent of the differences of the stabilities of the corresponding states. However, the results in Ref. 25 were limited in scope and had no clear physical explanation. The generalization of these conclusions to the other quantities is not obvious.

In this work, we use a chemical kinetic formalism to determine how steady-state probabilities and the flux distribution of the underlying biochemical reactions are affected by perturbations of the underlying free energy landscape. We illustrate the generality and the biological implications of our results by specifically studying three diverse biological systems. The examples include: (i) two alternative pathways for protein folding in the presence of misfolded error states, which is motivated by folding pathways in hen egg-white lysozyme^26,27^, (ii) a Hopfield KPR network describing aminoacyl(aa)-tRNA selection during protein translation in the *Escherichia coli* ribosome^19,25,28–31^, and (iii) the myosin-V motor protein that walks along actin cytoskeleton filaments^22,24^.

The results demonstrate that an arbitrary ratio of the steady-state fluxes is invariant to the energy perturbations of the stabilities of the discrete states, and therefore, is only dependent on the free energy barriers. In other words, we demonstrate that the biological properties that are expressed in terms of the steady-state flux ratios are governed by kinetic and not by thermodynamic factors. These results have wide-ranging implications for the types of genetic or chemical perturbations capable of changing the steady-state flux distribution through biochemical reaction networks. Generally, the values of the free energy barriers correlate with the free energies of the states^32^. However, even though mutations can perturb both the free-energy barriers and the stability of the minima, it is important to know that the latter energy parameters do not affect the flux ratios. Thus, our result can direct experimental measurements of the underlying free energy landscape.

## METHODS

### Notation and Setup

Consider an arbitrary biochemical system described by the linear formalism^12^, i.e., by a chemical kinetic network with quasi-first order transitions between *N* biochemical states (see Fig. 1*A*) with the rate constant for a reaction *i → j* (i.e., a transition from the *i*-th state to the *j*-th state) denoted as *k_i,j_*. Generally, there could be multiple elementary reactions between the states *i* and *j*. Therefore,

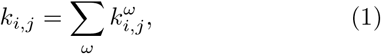

where the index *ω* runs over all of the possible reaction pathways *i → j*. For each elementary reaction the rate constants 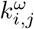 can be expressed as the product of a prefactor and an exponential free energy barrier term (e.g., as in transition-state theory):

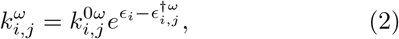

in which the prefactors of the bimolecular reactions that were initially second order (i.e., with cofactor binding steps) included the dependence on the chemical potentials *μ_γ_* of the these cofactors (see the SI for discussion and definition of the prefactor 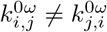). Here, *ϵ_i_* is the energy of the *i*-th state and 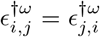 are the transition energies in units of *k_B_T* (see Fig. 1*B*).

**FIG. 1.**
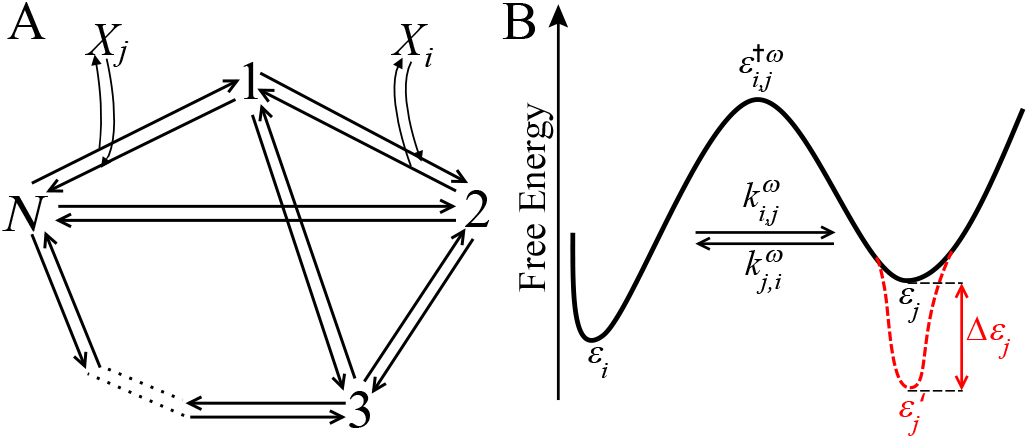
General biochemical kinetic network and its free energy landscape. (*A*) Biochemical network comprised of *N* different states. The discrete states are denoted by numbers 1, .., *N*, and the cofactor molecules in the bath have chemical identities *X_i_* and *X_j_*, respectively. (*B*) Free energy landscape for a reaction between the states *i* and *j* with well-depths *ϵ_i_* and *ϵ_j_* and the transition state energy 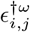. The rate constants 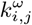 and 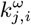 correspond to the forward and reverse reactions between the two states *i* and *j* on the pathway *ω*. Note that the energy of the state *j* is decreased by an amount Δ*ϵ_i_* (red, dashed curve) such that its energy is 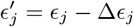.

In the limit of *t → ∞*, the system is characterized by a stationary probability distribution vector with *N* components, **P** = [*P*_1_, *P*_2_…, *P_N_*]^*T*^ that can be obtained by solving a set of steady-state equations subject to the normalization condition:

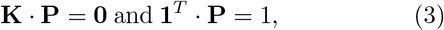

where **1** is an *N* × 1 unit vector, and the **K** is an *N × N* rate matrix:

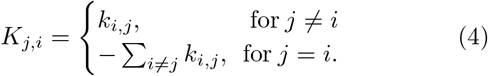

Assuming this matrix is not rank deficient, (3) has a unique solution for **P**. The corresponding steady-state fluxes for *i* → *j* are given by 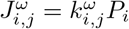

Depending on whether the rate constants 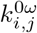 are subject to the detailed balance constraints, this formalism can describe both equilibrium and non-equilibrium steady states. In the former case, the fluxes balance 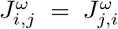 and the stationary probability distribution corresponds to the Boltzmann distribution^15^. Below we explore non-equilibrium steady-state fluxes and the probabilities.

### Perturbation of the Free Energy Landscape

Consider a perturbation that decreases the energy of an arbitrary state *m* on the free-energy landscape of the system by an amount Δ*ϵ*_*m*_, i.e., such that its energy changes to 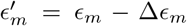 (see Fig. 1*B*). Here and below the prime symbol ′ denotes the parameters following this perturbation. Assume that the energies of other states, namely, the transition state energies 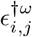 and the chemical potentials *μ_γ_* [see SI Eqs. (S8-S10)] of all of the cofactor molecules in the bath are held constant.

From (2), the perturbation reduces all of the rate constants for the reactions *m* → *j* by a factor of 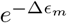, but all of the other rate constants 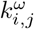 remain the same. As a result, the perturbed steady-state probability distribution **P**′ satisfies equations of the form of (3) with the perturbed matrix **K**′:

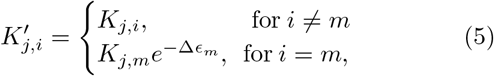

i.e., with the entire *m*-th column scaled by a factor of 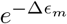. It is not hard to show [see the SI Eq. (S15)] that the perturbed probabilities 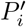 can be expressed in terms of the unperturbed probabilities *P_i_* as follows

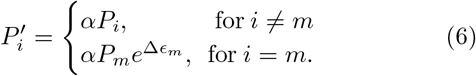

with normalization constant 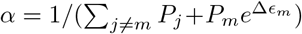 that guarantees the conservation of probability, **1**^*T*^ · **P**′ = 1.

### Invariance of the Ratios of the Steady-State Fluxes

From (6), we conclude that all of the steady-state fluxes are scaled by the same factor:

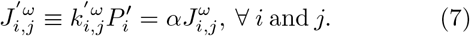

The factor *α* cancels out for all of the ratios of the perturbed steady-state fluxes, and we demonstrated that

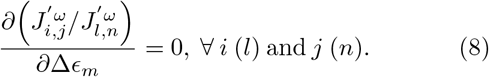

Therefore, we conclude that *the ratios of stationary fluxes or any of their linear combinations do not depend on the energies of the individual states ϵ_i_ and only depend on barrier heights* 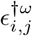.

### Illustrative Examples

To illustrate the implications of (8), we use three major biological processes: protein folding networks inspired by hen egg-white lysozyme, a KPR network for aa-tRNA selection during protein translation in the *E. Coli* ribosome, and the myosin-V motor protein that walks on the actin cytoskeleton filaments. For simplicity of the notation, we drop all of the prime ′ and *ω* superscripts on the perturbed steady-state fluxes.

### Data Availability

The kinetic parameters used for illustrative examples from Refs.^24,27,30,31,33,34^ are given in the SI Tabs. S1-S3.

## RESULTS

## RESULTS

### Protein Folding Network

Protein folding is a fundamental process that is required for all living cells to maintain their proper functionality. Indeed, misfolded proteins and their aggregation have been linked to many types of pathological diseases^35^. Protein folding occurs via a series of different funnels on a free energy landscape, and as such, the process can be described using the chemical kinetic formalism for biochemical networks^15,26,27^. Protein folding often does not require any energy consumption, and therefore, it can be studied within the equilibrium thermodynamics framework^15,35^. On the other hand, in live cells all proteins are synthesized in the unfolded states and degraded/diluted in the relatively stable folded states. In this situation, a non-equilibrium steady-state distribution of fluxes in a protein folding network can give information about the relative importance of different folding pathways and the probability of reaching incorrect metastable folding states.

Here, we consider a kinetic scheme representing the protein folding inspired by hen egg-white lysozyme^27,33^. For this enzyme, two mechanistic descriptions, namely, independent unrelated pathways (IUP) and pre-determined pathways with optional error (PPOE), have been proposed. The IUP model (also known as a heterogeneous folding) assumes that protein folding occurs through various intermediate states on different pathways that have no relationship to one another^26^. On the other hand, the PPOE model supposes that all proteins fold cooperatively through the same productive pathway via partially folded subunits called foldons that resemble the correctly folded structure. However, during the folding process, the protein may also become diverted into multiple misfolded error states that hinder productive folding^26^.

We chose a kinetic scheme (see Fig. 2*A*) that combines the physicochemical aspects of both the IUP and PPOE models. There are two possible pathways for the unfolded protein in state 1 (U) to reach the folded (native) conformation in state 3 (F) on this biochemical kinetic network. This means that productive protein folding can proceed independently through either of the intermediate states 2 or 4. However, the protein may also become trapped in the misfolded error states denoted respectively as 5 and 6 on the two folding pathways. The misfolded error states 5 and 6 can be viewed as traps on the free-energy landscape that might kinetically block productive protein folding into the folded conformation (state 3) on the two different pathways. To realize the non-equilibrium steady state and the non-zero fluxes in this network, we assume that the protein is synthesized in state 1 and degraded from state 3 with the rate constant *k*_⊘,1_. In the non-equilibrium steady state, the synthesis and degradation fluxes are balanced so that we can map this scheme to the formalism considered in the Methods by effectively introducing an irreversible 3 → 1 transition with the rate *k*_⊘,1_ (similar to Ref. 36).

**FIG. 2.**
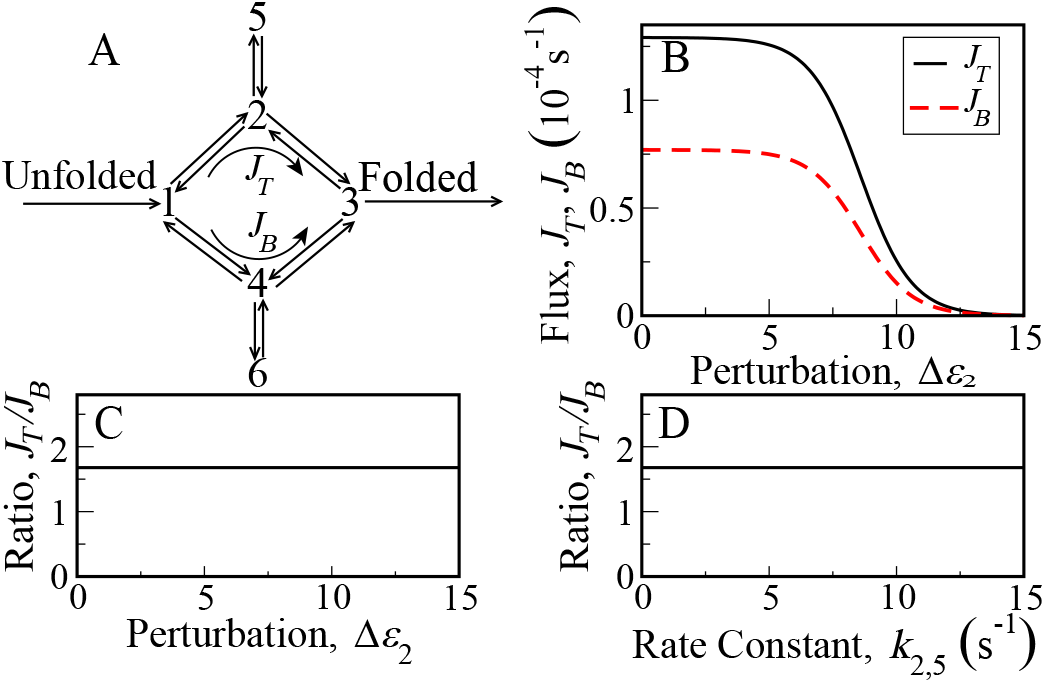
Steady-state fluxes and their ratio for a protein folding network from folding pathways in hen egg-white lysozyme from Refs. 27,33. (*A*) Protein folding network through two independent pathways via intermediate states 2 and 4 (with misfolded error states 5 and 6) to reach the folded conformation in state 3 from the unfolded state 1. (*B*) Steady-state fluxes *J_T_* and *J_B_* that decay to zero as a function of the energy perturbation Δ*ϵ*_2_. (*C*) Ratio of the steady-state fluxes *J_T_/J_B_* as a function of the energy perturbation Δ*ϵ*_2_. (*D*) *J_T_/J_B_* as a function of the rate constant *k*_2,5_ for the reaction 2 → 5.

We can now apply our analysis to ask how the steady-state folding flux splits between the top pathway via state 2 (*J_T_* = *k*_1,2_*P*_1_ − *k*_2,1_*P*_2_) and the bottom pathway via state 4 (*J_B_* = *k*_1,4_*P*_1_ − *k*_4,1_*P*_4_). (8) implies that the steady-state flux ratio will be independent of the stability of the intermediate states and is only affected by the free energy barriers. For example, consider a change in the energy of the state 2 by Δ*ϵ*_2_. This perturbation will change both *J_T_* and *J_B_* proportionally in Fig. 2*B*. As a result, the ratio of the steady-state fluxes *J_T_*/*J_B_* does not change as a function of Δ*ϵ*_2_ (see Fig. 2*C*). The result is also apparent from the analytic expression for *J_T_*/*J_B_* [see the SI Eq. (S27)]. Notably, the presence of the misfolded error states 5 and 6 is mathematically equivalent to the perturbations of the energy of the states 2 and 4, respectively. Therefore, the ratio of the steady-state fluxes *J_T_ /J_B_* does not depend on the rate constant going into or out of these states (e.g., *k*_2,5_ in Fig. 2*D*). Thus, the stability or presence of a misfolded error state does not affect how the steady-state flux splits between the two folding pathways. Notably, identical results can also be obtained using the approach developed in Ref. 36. However, our approach is arguably more general as it extends to the networks with futile cycles.

### KPR Network

Enzymatic catalysis in the presence of right (cognate) and wrong (non-cognate) substrates can often achieve high selectivity through KPR mechanisms with the use of GTP/ATP hydrolysis energy. For such networks, the flux distribution can be used to quantify the probability of reaching the wrong product state (error rate) or the probability of futile cycles that result in hydrolysis without any product formation. Here we consider a KPR network that enhances the accuracy of aa-tRNA selection in the *E. Coli* ribosome^19,28,29,34^ during protein translation, and it is shown in Fig. 3*A*. It contains two symmetric cycles involving the right R (cognate) or the wrong W (noncognate) aa-tRNA molecules and two ways to reset from states 3/5 to the initial state 1. This can occur via a catalytic step resulting in product formation (rates 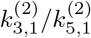) or via a proofreading step that results in aa-tRNA excision (rates 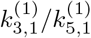). The steady-state ratios of the corresponding fluxes define important system properties of the enzyme. The error rate can be defined as the fraction of times the non-cognate amino acid is incorporated into a protein, i.e., as a ratio of 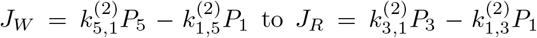. Perturbations of the energy of an intermediate state (e.g., state 3 in Fig. 3*B – D*) will change the individual fluxes in Fig. 3*B*, but the ratio of the fluxes in Fig. 3*C* will remain invariant. This is a straightforward consequence of (8) and it generalizes the result of Ref.^25^. We can also conclude that the error rate will be invariant to perturbations of the discrete-state energies even in the presence of multiple competing pathways, multiple intermediate states, and/or multiple proofreading reactions. Therefore, the error rate in more complex KPR networks is under kinetic control as long as it can be expressed as a ratio of the steady-state fluxes in the linear chemical kinetic framework.

**FIG. 3.**
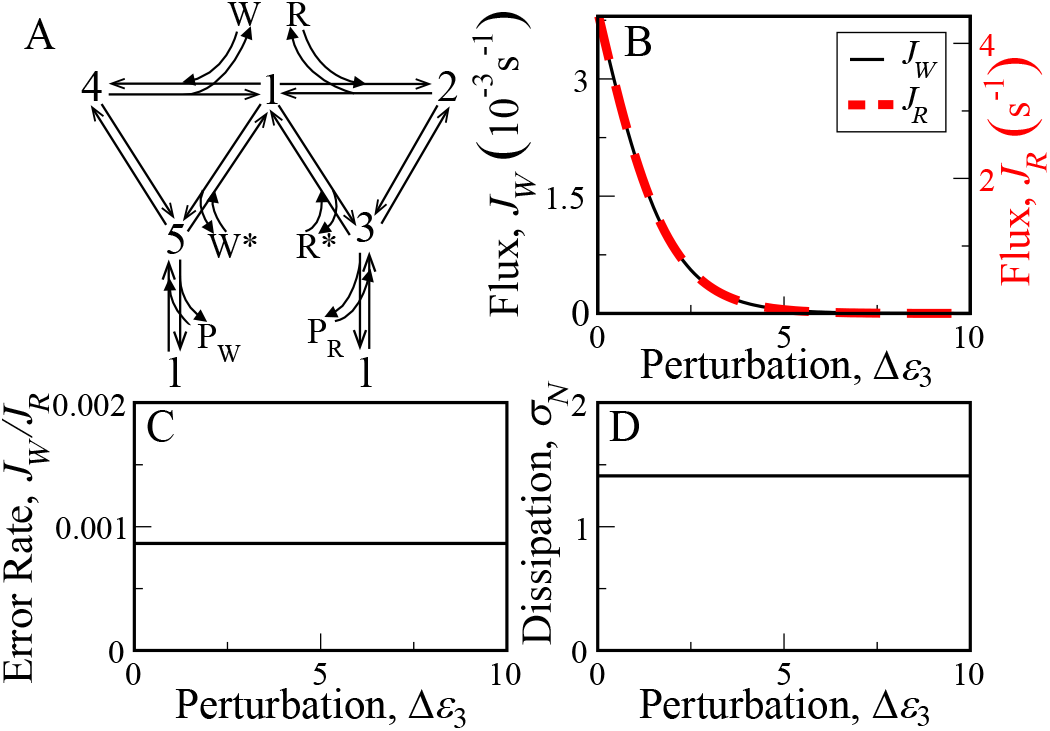
Steady-state fluxes and their ratios for the KPR network describing aa-tRNA selection in the *E. Coli* ribosome. (*A*) KPR scheme where the right R or the wrong W aa-tRNA molecule can bind to the free enzyme state 1. The network from kinetic schemes in Refs. 25,30,31 is comprised of two cycles: a proofreading cycle with R* or W* and a product formation cycle with products P_R_ or P_W_. (*B*) Steady-state fluxes for addition of the wrong amino acid *J_W_* and the right amino acid *J_R_* that decay to zero as a function of the energy perturbation Δ*ϵ*_3_. (*C*) Error rate *J_W_*/*J_R_*. (*D*) Normalized energy dissipation *σ_N_*.

Energy efficiency of the ribosome can be quantified by looking at the number of GTP molecules hydrolyzed per peptide bond^19^ or as the energy dissipation per product formed normalized by the GTP hydrolysis energy, i.e.,

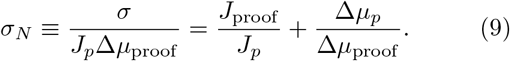

Here, *σ* is the total energy dissipation in the non-equilibrium system, while Δ*μ*_proof_ and Δ*μ_p_* are the respective chemical potential differences of the proofreading and the catalytic cycles in the biochemical network. The product formation flux is *J_p_* = *J_R_* + *J_W_*, and the proofreading flux is given by 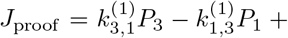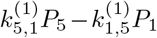. The ratio of these fluxes and the normalized energy dissipation *σ_N_* will again be invariant to the energy perturbation of an intermediate state, e.g., Δϵ_3_ (see Fig. 3*D*).

### Myosin-V Network

Motor proteins belong to a broad class of enzymes that convert chemical energy (e.g., from ATP hydrolysis) to perform mechanical work in cells. For example, motor proteins in cells that hydrolyze ATP use the hydrolysis energy to walk on the cytoskeletal track while transporting loads of intracellular cargo^20,22^. The motors are often described by kinetic models^22^ that allow one to predict the flux distribution that defines important properties such as the probability of forward or backward steps or probability that the motor proteins hydrolyze ATP but remain at the same position on the track.

Here, for illustration we examine a kinetic model for myosin-V in Fig. 4*A* that shows the intermediate states through which the motor protein proceeds as it attempts to move one ~ 36 nm step along an actin filament^24^. The myosin-V network has a main stepping cycle (via states 1-5) where the motor protein takes one step forward or backward on the the actin filament and a futile cycle where the motor protein makes no net movement on the actin filament (via states 2, 3, 4 and 6). During the futile cycle, myosin-V does not move one step along the actin filament and all of the energy from ATP hydrolysis is wasted. This is not the case for the main stepping cycle, where myosin-V uses the energy available from ATP hydrolysis to take one step forward along the actin filament while doing mechanical work against a load.

**FIG. 4.**
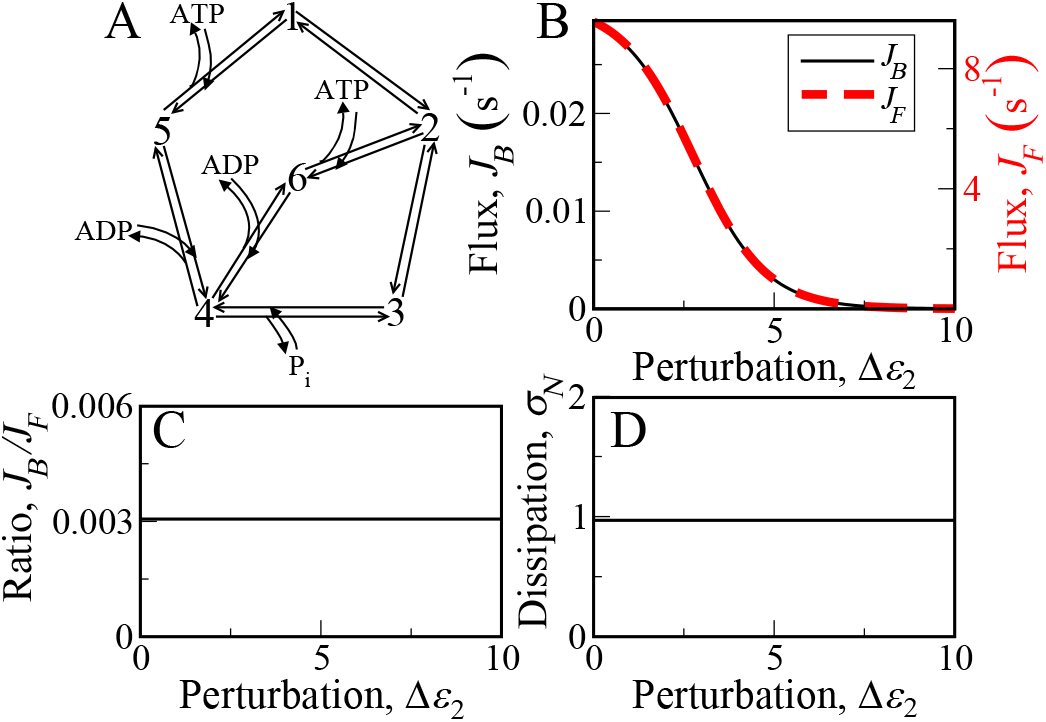
Steady-state fluxes and their ratios for the mechanochemical network of the two-headed myosin-V motor protein while taking one step on an actin cytoskeleton filament. (*A*) Mechanochemical network for the myosin-V motor protein from Ref. 24 comprised of two cycles: a futile cycle and a main stepping cycle. (*B*) Steady-state fluxes for backward steps *J_B_* and forward steps *J_F_* of myosin-V that decay to zero as a function of the energy perturbation Δ*ϵ*_2_. (*C*) Ratio of backward to forward steps *J_B_*/*J_F_*. (*D*) Normalized energy dissipation *σ_N_*.

The fraction of times that myosin-V takes a backward step is given by the ratio of the backward stepping flux *J_B_* = *k*_2,1_*P*_2_ to the forward stepping flux *J_F_* = *k*_1,2_*P*_1_. If the energy of an intermediate state is perturbed, e.g., by Δϵ_2_, the individual fluxes *J_B_* and *J_F_* in Fig. 4*B* will change as well. However, the ratio of the backward to forward fluxes in Fig. 4*C* is invariant to the energy perturbation. This result is easily seen to be a consequence of (8) for the motion of the myosin-V motor protein. Moreover, we can conclude that this ratio of the backward and forward stepping fluxes will remain invariant regardless of how many intermediate states or futile cycles are present on the mechanochemical network.

The energy efficiency of myosin-V can be defined as the fraction of ATP hydrolysis energy consumed during one step on the track^22^ or as the energy dissipation per step normalized by the energy expended from ATP hydrolysis, i.e.,

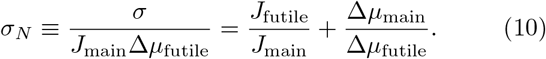

Here, *σ* is the total energy consumed during the mechanochemical process and Δ*μ*_futile_ and Δ*μ*_main_ are the respective chemical potential differences of the futile cycle and the main stepping cycle. The main stepping flux is *J*_main_ = *k*_1,2_*P*_1_ − *k*_2,1_*P*_2_, and the futile flux is *J*_futile_ = *k*_6,2_*P*_6_ − *k*_2,6_*P*_2_. The ratio of these two fluxes and the normalized energy dissipation *σ_N_* are also invariant to energy perturbations of an intermediate state such as *Δϵ*_2_ on the mechanochemical cycle (see Fig. 4*D*).

## DISCUSSION

Our main theoretical result shows that the ratios of steady-state fluxes are independent of energy perturbations of the intermediate states on biochemical networks that describe a wide class of biological processes. This result can be explained using the following arguments. The energy perturbation at the specific site leads to a change in the stationary probability of the perturbed state and the transition rates out of the perturbed state. However, the effect of the perturbation is opposite for these two properties. If the perturbation increases (decreases) the stationary probability by the exponent of the energy perturbation, it simultaneously decreases (increases) the outgoing transition rates by the same factor. In addition, because the total probability in the system is conserved, the stationary probabilities of all of the states should be modified by the same normalization parameter. This leads to changes in all of the steady-state fluxes in the system by the same parameter, including the fluxes out of the perturbed state where the changes in the stationary probability is compensated by changes in the outgoing transition rates. As a result, all of the steady-state fluxes are scaled in the same way and their ratios remain constant.

The important implication of our theoretical results is that the properties of major biological processes that depend on the ratios of steady-state fluxes are governed by kinetic rather than thermodynamic (energetic) factors of the underlying free energy landscapes. Specifically, the ratios of any two steady-state fluxes depend on the transition state energies, but *not* on the energies of the discrete (internal) states on the free-energy landscape. Consequently, the physicochemical properties of biological systems that depend on the ratios of the steady-state fluxes are exclusively under kinetic control, and any changes in the well-depths of the individual states do not affect these properties. Therefore, the only way to modify these properties is to change the energy barriers between the intermediate states. Moreover, the invariance result is still valid if several energy perturbations are simultaneously occurring in the system because these perturbations are local and the total effect of the perturbations is additive. However, the properties of the systems expressed in terms of individual fluxes and/or state probabilities are still affected by perturbations of the discrete-state energies and therefore, are subject to a combination of thermodynamic and kinetic control.

Our theory is applicable to kinetic models of biochemical systems that, from a single-molecule perspective, are described by linear master equations, i.e., they only involve quasi-first-order transitions between the discrete states. Despite this limitation, the framework still applies to a wide-range of biological processes under the commonly used assumption of time-scale separation^12^. These processes include enzyme catalysis and allosteric control, processive motion of molecular motors, receptor signal-transduction, ion-channel transport, transcriptional regulation, and post-translational modification^12^. To illustrate the biological relevance of our theoretical findings, we applied them to three major biological processes, namely, protein folding via multiple pathways, aa-tRNA selection during protein translation, and the myosin-V molecular motor dynamics. Our examples reveal some counter-intuitive implications for specific experimentally observable quantities.

For the protein folding network motivated by the hen egg-white lysozyme system, we find that the splitting fractions for the different pathways along which the protein can fold do not depend on the energy (well-depths) of the intermediate states as long as the transition-state energies are constant. Mathematically, such perturbations are equivalent to the introduction of misfolded error states^26,27^. Our invariance results imply that the existence of such states or generally the existence of multistate dead-end branches from the intermediate states will not affect the splitting fractions. The results are clearly generalizable to kinetic models of protein folding in which multiple stable folding conformations are present, e.g., when there is a possibility of prion states^37^. Our results also imply that the flux splitting between the native and prion conformations would require changes in the free energy barriers and not in the well-depths along each of the folding pathways. Thus, a drug that binds to and stabilizes the intermediate but needs to dissociate before the transition states are reached cannot affect the flux splitting.

Our examination of the biochemical network for aa-tRNA selection in the *E. Coli* ribosome^19,23,25,30,31^ demonstrates that our invariance statement holds in the presence of energy-dissipating cycles on the network. We show that both the error fraction and the normalized energy dissipation can be expressed as flux ratios, and therefore, they are invariant to changes in the energies of the intermediate states on the underlying free-energy landscape. Hence, genetic mutations in the ribosome^28,34^ that cause changes in these properties must affect the kinetic features of the free energy landscape.

Finally, for the physicochemical properties of myosin-V we show that the ratio of the forward and backward stepping fluxes and the normalized energy dissipation are invariant to energy perturbations of all of the intermediate states. This indicates that the energy expenditure from ATP hydrolysis during the processive motion of the myosin-V motor protein^20,22,24^ is exclusively under kinetic control as long as the external force *F*_external_ remains constant. Consequently, drugs that specifically stabilize different states on the mechanochemical network for myosin-V but have to dissociate before the transition to the next or to the previous state can occur do not affect the ratio of the energy efficiency or the ratio of the forward to backward steps.

We note that the stationary flux ratios will change if the energy perturbations of individual states are coupled to changes in the free energy barriers. However, our invariance results can have important implications for such cases. Identical effects on the steady-state flux ratios are expected from two mutations stabilizing the transition state by the same mechanism and thus, resulting in identical changes of the transition state energy barriers 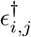. That will be the case even if mutations affect the state stabilities *ϵ_i_* differently (see SI discussion on energy coupling in protein folding from Φ-value analysis). Therefore, even for the perturbations affecting both the energy barriers and the energies of the states, it is important to know that the changes in the latter do not affect the physical properties that depend on the flux ratios.

Our theoretical study presents a general statement on how kinetic models that provide a quantitative description of a wide class of biological systems can be affected by the underlying features of the free-energy landscape. This result pinpoints a hidden symmetry in the underlying equations, which places important constraints on the mechanism for the regulation of important system-level properties. It will be important to see the implication of the invariance results for other biological systems and to determine if the result can be extended to an asymptotic flux behavior away from the steady state.

## ACKNOWLEDGEMENTS

This work was supported by Center for Theoretical Biological Physics National Science Foundation (NSF) Grant PHY-1427654. A.B.K. also acknowledges support from the Welch Foundation Grant C-1559 and from NSF Grant CHE-1664218. O.A.I. also acknowledges support from the Welch Foundation Grant C-1995. We thank Qiwei Yu for useful suggestions.

## AUTHOR CONTRIBUTIONS

A.B.K. and O.A.I. designed research; J.D.M., A.B.K., and O.A.I. performed research; J.D.M., A.B.K., and wrote the paper. *O.A.I. and A.B.K. contributed equally to this work. To whom correspondence should be addressed.

### SIGNIFICANCE STATEMENT

Chemical kinetic formalism is widely used to describe biochemical processes, including channel transport, enzymatic catalysis, genetic regulation, and signal transduction. Fundamentally, transitions between biochemical states are defined by the underlying free-energy landscapes, but few general results relate the landscape features to biologically relevant properties. At equilibrium, the probabilities of discrete states only depend on their energies and not on the energy barriers in-between states. However, few general claims about non-equilibrium steady states have been formulated. Here, for non-equilibrium steady states of networks with quasi-order transitions, we prove that the flux ratios are invariant to energy perturbations of the states, and therefore, are only affected by the free-energy barriers. Biological implications of the result are illustrated for three distinct biological processes.

## Supporting Information

### Invariance Proof for an Arbitrary Biochemical Kinetic Network

#### Notation and Setup

Consider an arbitrary biochemical kinetic network with quasi-first order transitions between *N* biochemical states (see Fig. S1*A*) that is described by the linear chemical kinetic formalism^1^. The rate constant for the reaction *i* → *j* (i.e., a transition from the *i*-th to *j*-th state with *i ≠ j*) is denoted by *k_i,j_*. Note that in general there are multiple elementary reactions between the two states *i* and *j* such that the rate constant *k_i,j_* can be decomposed leading to

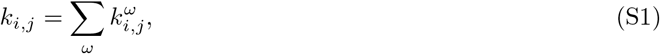

where the index *ω* runs over all of the possible reactions *i → j*. The occupancy probability vector **P**(*t*) has *N* components, which are the time-dependent probabilities of finding the system in a given state at time *t*,

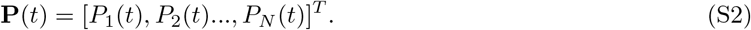

The system of forward master equations that governs the temporal evolution of the system is solved to obtain the normalized probability distribution **P**(*t*),

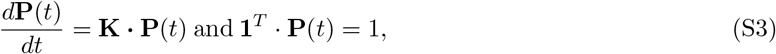

where **1** = [1,…, 1]^*T*^ is a unit *N* × 1 vector, and the elements of the *j*-th row and the *i*-th column of the *N* × *N* rate matrix **K** are given by

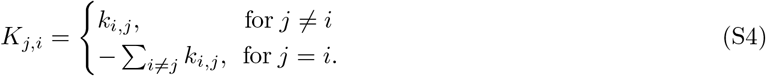

**Figure S1.**
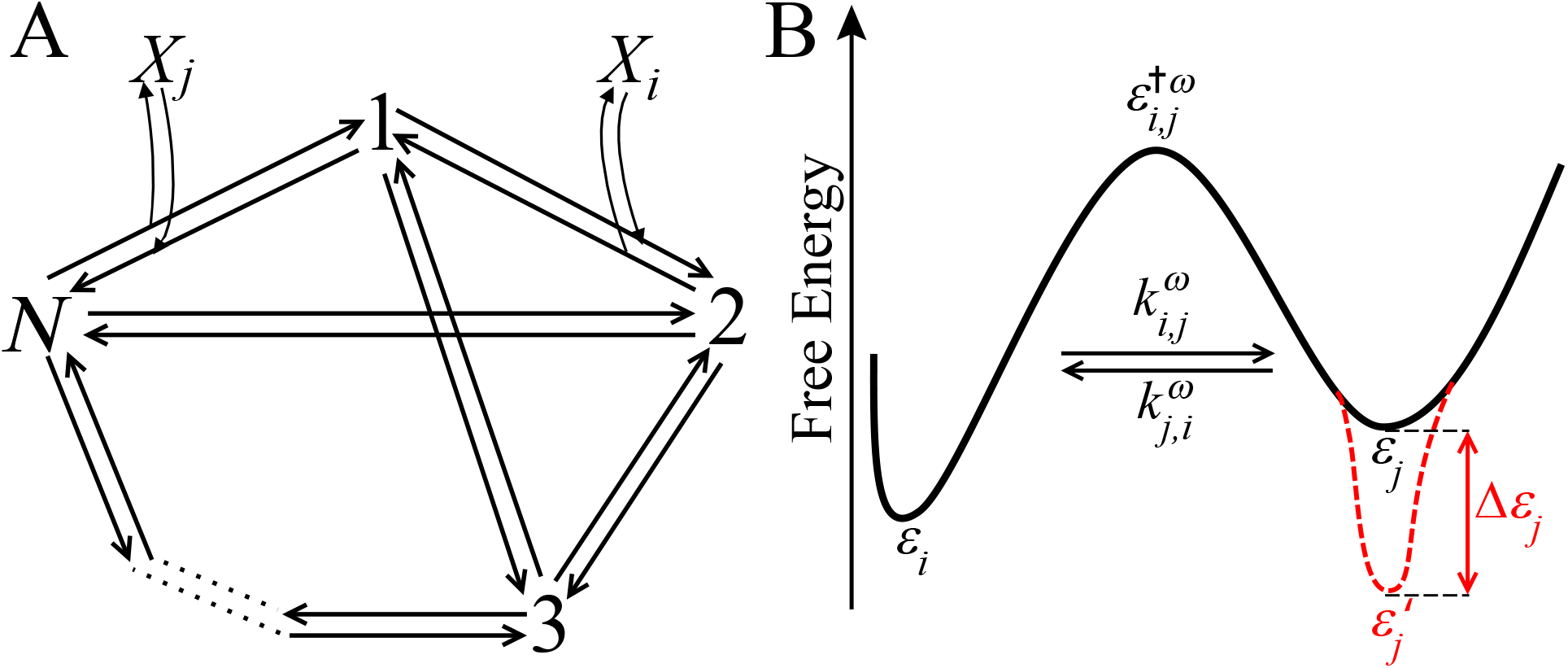
Representation of a general biochemical kinetic network and a portion of its free energy landscape. (*A*) General biochemical network comprised of *N* different states. The various internal states of the system are denoted by the numbers 1, .., *N*, while the cofactor molecules in the bath that are involved in the various bimolecular reactions have chemical identities denoted by *X_i_* and *X_j_*, respectively. (*B*) A portion of a free energy landscape corresponding to the reaction between the states *i* and *j* with the energies of the respective intermediate states (well-depths) denoted by *ϵ_i_* and *ϵ_j_* and the transition state energy denoted by 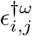. The respective rate constants 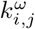 and 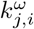 correspond to the forward and reverse reactions between the two states *i* and *j* on the pathway *ω*. Note that the energy of the state *j* is decreased (perturbed) by an amount Δ*ϵ_j_* (red, dashed curve) such that the energy of the perturbed state is 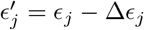.

In the limit of the time *t* → ∞, the system approaches a non-equilibrium steady-state denoted by the stationary occupancy probability vector **P** such that

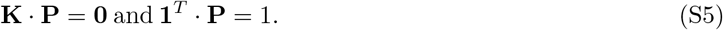

Assuming that the matrix **K** is not rank deficient, these equations define a unique solution to the set of steady-state forward master equations, i.e., the stationary probability distribution **P**. Note that the stationary probability distribution **P** would be the Boltzmann distribution if the system approached thermodynamic equilibrium in the limit of *t* → ∞. In this case, the probability of the individual states is known immediately as 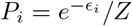 where *ϵ_i_* is the energy of the state and 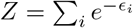 is the partition function from statistical mechanics^2^.

#### Elementary Reactions and the Free Energy Landscape

Consider the arbitrary biochemical kinetic network in Fig. S1*A* with *N* internal states that define the system contained within a bath (i.e., the cellular environment) of chemical species defined as cofactor molecules with chemical identities *X_γ_*. The steady-state of the system defined in Eq. (S5) will change in response to chemical reactions occurring between the internal states that affect the energies *ϵ_γ_* (in units of *k_B_T*) of the cofactor molecules in the bath. The rate constants 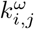 corresponding to the reactions *i* → *j* in the system such that they can be decomposed and expressed as

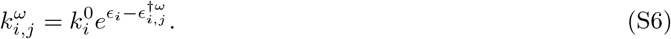

Here, 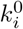 is a pre-exponential (frequency) factor independent of the energies of the internal states in the system, while *ϵ_i_* and 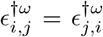 are the respective energies of the internal and the transition states on the free energy landscape.

However, changes in the energies Δ*ϵ_γ_* of cofactor molecules that affect the chemical composition of the bath due their participation during elementary reaction steps (i.e., binding or dissociation) must be taken into account in our theory. To this end, any rate constants 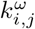 for bimolecular and higher order reactions in the system that cause changes in the energy *ϵ_γ_* of the cofactor molecule *X_γ_* due to a binding reaction are modified by a prefactor 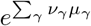. The prefactor effectively raises the energy *ϵ_i_* of the internal state and reduces the barrier for the reaction by ∑*_γ_ν_γ_μ_γ_* Specifically, *ν_γ_* is the stochiometric coefficient denoting the number of cofactor molecules *X_γ_* that are consumed during the binding reaction. Note that *ν_γ_* = 0 if no cofactor molecule *X_γ_* participates in the reaction such that the elementary reaction step does not change the chemical composition of the bath. Moreover, *μ_γ_* ≡ Δ*ϵ_γ_* is the fixed chemical potential defined as the energy change per cofactor molecule *X_γ_* during a binding reaction that consumes this species ^3^. Note also that if the biological process involves a mechanochemical cycle for a motor protein, the rate constant is modified by a prefactor of 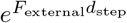 for the elementary reaction where the mechanical work is performed. In this case, *F*_external_ is the opposing force due to the external load from the intracellular cargo and *d*_step_ is the distance that the motor protein moves in one step on the cytoskeleton track ^4^.

The modification of the unimolecular prefactors 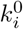 for the binding reactions of the cofactor molecules *X_γ_* enables the rate constants to be written as

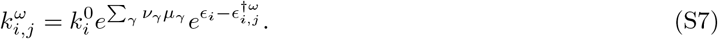

The prefactor 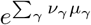 is treated as a constant because it corresponds to changes in the energies of the cofactor molecules in the bath and not to changes in the energies of the internal states of the system. Therefore, a new set of pre-exponential factors 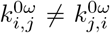 can be defined with the prefactor absorbed into them,

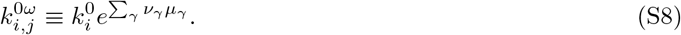

This leads to the following expression for all of the rate constants 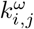 of the elementary reactions on the biochemical network,

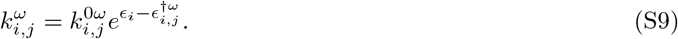

#### Generalized Detailed Balance Constraints

Consider an arbitrary cycle on a biochemical kinetic network with a chemical potential difference (i.e., a thermodynamic force) Δ*μ*_cycle_. The generalized detailed balance constraint in terms of the rate constants for the clockwise 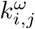 and counterclockwise 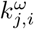 directions of the cycle is given by

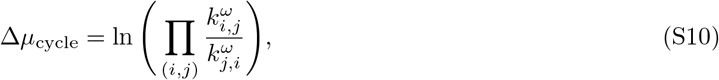

where the product runs over all pairs of transitions (*i, j*) connected on the cycle. Using Eq. (S7) for the rate constants 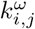 in this equation, it is easy to see that Δ*μ*_cycle_ defines the chemical potential difference between all of the cofactor *X_α_* (substrate) and *X_β_* (product) molecules that participate in the binding and dissociation reactions of the cycle,

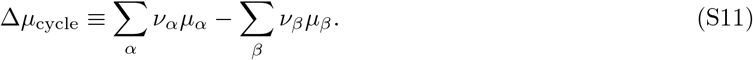

Note also that using the expression for the rate constants from Eq. (S9) in Eq. (S10) leads to the following equation for the chemical potential difference,

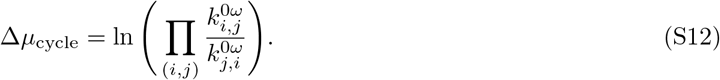

Substitution of the prefactors 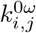 defined in Eq. (S8) into this equation gives the defintion for the chemical potential difference over the cycle in Eq. (S11). Therefore, the generalized detailed balance constraint for the cycles of biochemical networks arises naturally as a consequence of the linear chemical kinetic formalism.

If the cycle does not change the chemical composition of the bath, it obeys the detailed balance constraint such that the chemical potential difference over the cycle Δ*μ*_cycle_ = 0. This implies that when the cycle runs, e.g., in the clockwise direction, binding of a cofactor molecule *X_α_* is compensated by the dissociation of the same molecule *X_β_* = *X_α_* such that the species is not chemically modified upon completion of the cycle. If the cycle runs in the counterclockwise (i.e., the reverse) direction, the dissociation of the cofactor molecule *X_β_* = *X_α_* would be a binding reaction and visa-versa. In this scenario, the prefactors cancel out because they are the same (i.e., 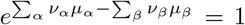) regardless of whether the cycle proceeds in the clockwise or counterclockwise directions. Therefore, cycles at thermodynamic equilibrium do not change the chemical composition of the cofactor molecules in the bath. However, if the cycle does change the chemical composition of the bath, the chemical potential difference over the cycle Δ*μ*_cycle_ ≠ 0, and the cycle violates the detailed balance constraint. In this situation, binding of the cofactor molecule *X_α_* results in the production of a new cofactor molecule *X_β_* upon completion of the cycle. Thus, the respective prefactors for the clockwise and counterclockwise directions of the cycle are not the same, and the cycle affects the concentrations of the species [*X_α_*] and [*X_β_*] in the bath. For dilute solutions of cofactor molecules (e.g., *X_α_*) the chemical potential depends explicitly on the concentration [*X_α_*],

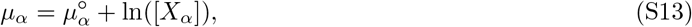

where 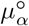 is the chemical potential of the cofactor molecules *X_α_* when the concentration of this species is 1 M. Here, it is clear that the prefactor for the binding reaction is proportional to 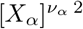. Using Eq. (S13) in Eq. (S11), the chemical potential difference over the cycle is given by

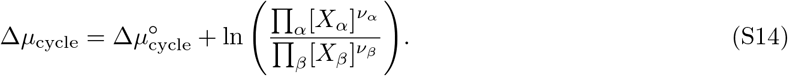

For example, the production of the new cofactor molecule *X_β_* during the cycle may lead to changes in the concentrations of other energy-rich molecules such as [ATP] and its hydrolysis products [ADP] and [P_i_] such that the chemical potential difference for ATP hydrolysis is given by

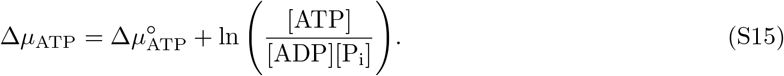

Consequently, the composition of the bath changes for a cycle in a non-equilibrium steady-state due to chemical modification of the cofactor molecule *X_α_* to *X_β_* and/or energy expenditure from ATP hydrolysis. This result demonstrates that the expressions for the rate constants in the linear chemical kinetic formalism provide a way to properly account for the detailed balance constraints for the cycles on biochemical networks.

#### Perturbation of the Free Energy Landscape

Consider a perturbation that decreases the energy of an arbitrary state *m* on the free energy landscape of the system by an amount Δ*ϵ_m_* (in units of *k_B_T*) such that 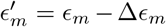, while all of the energies of the other intermediate states *i* ≠ *m* remain unperturbed (see Fig. S1*B*). Moreover, all of the transition state energies 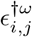 and the energies *ϵ_γ_* of all of the cofactor molecules are held constant. The perturbation of the state *m* reduces all of the rate constants 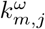 for the reactions *m* → *j* by a factor of 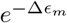, but all of the other rate constants 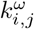 remain the same. Therefore, the rate constants 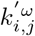 after energy perturbation of the state *m* are denoted by

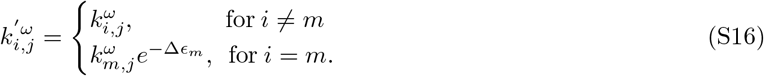

This leads to the scaling of the entire *m*-th column of the perturbed rate matrix **K**′ by the factor of 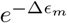 such that

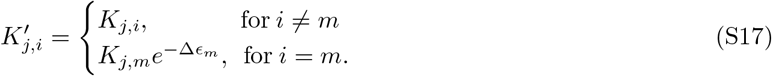

We can denote the new steady-state probability distribution as **P**′, and it must satisfy the following set of steady-state forward master equations and obey the conservation of probability,

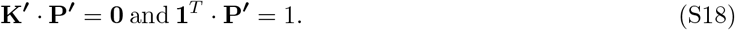

The perturbed probabilities can be expressed in terms of unperturbed probabilities according to

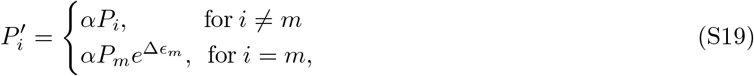

where *α* is a normalization constant,

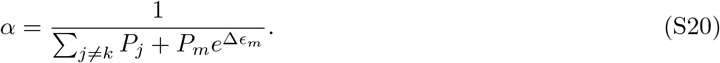

This choice of normalization constant guarantees that the conservation of probability is satisfied such that **1**^*T*^ · **P**′ = 1. Also, we must show that **K**′ · **P**′ = **0** by considering the *j*-th row of the vector equality

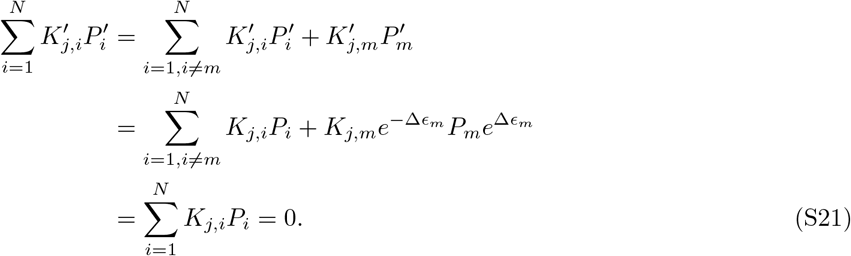

#### Invariance of the Ratios of the Steady-State Fluxes

By denoting 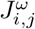 as the steady-state flux of the reaction *i* → *j*, we determine the relationship between the unperturbed steady-state flux 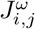 and the perturbed steady-state flux 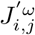. All of the unperturbed steady-state fluxes 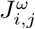 are scaled by the same factor *α* such that the perturbed steady-state fluxes 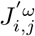 are given by

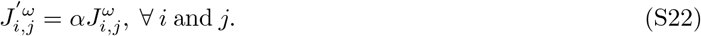

The unperturbed steady-state flux is 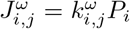, and the perturbed steady-state flux is 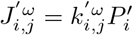 such that using the Eq. (S16) for 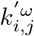 we obtain

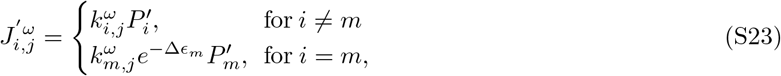

which can be expressed in terms of the unperturbed probabilities *P_i_* (see Eq. (S19)) as

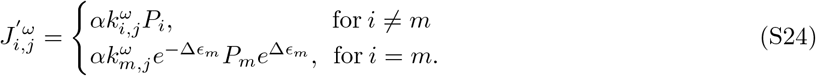

Due to the cancellation of the exponential terms involving the energy perturbation Δ*ϵ_m_*, we have shown that

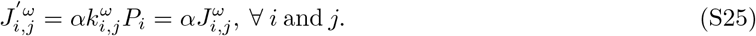

Note that the factor *α* cancels out for all of the ratios of the steady-state fluxes,

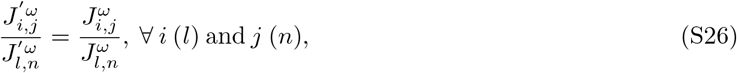

(and hence, for any linear combination of the fluxes) such that they are invariant to the energy perturbation Δ*ϵ_m_* of the *m*-th intermediate state on the free energy landscape. Therefore, we demonstrated that

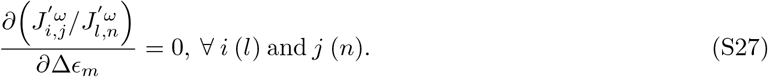

### Protein Folding Network

Table. S1 lists the kinetic parameters for the protein folding network motivated by folding pathways in the hen egg-white lysozyme ^5,6^ (see Fig. 2*A – D* in the main text). The unfolded protein in state 1 can reach the folded (native) conformation in state 3 along two possible pathways via the intermediate states 2 and 4. However, the protein may also become trapped in the misfolded error states 5 and 6, respectively. The correctly folded conformation (state 3) is degraded with a steady-state flux *J*_⊘,1_ = *k*_⊘,1_*P*_3_ such that there is an additional pathway 3 → 1 between the folded state 3 and the unfolded state 1. Consequently, the protein folding network is in a non-equilibrium steady-state where the folded proteins are degraded with a rate *k*_⊘,1_.

#### Kinetic Parameters: Protein Folding in Hen Egg-White Lysozyme

**Table S1:**
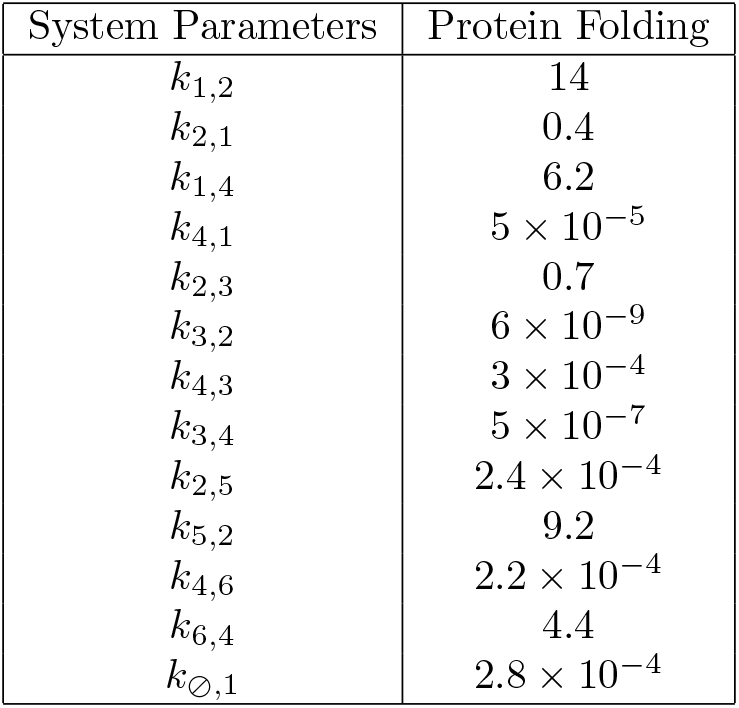
Rate Constants *k_i,j_* for Protein Folding motivated by Hen Egg-White Lysozyme.

The values of the rate constants *k_i,j_* (in units of s^−1^) for protein folding (except for *k*_2,5_, *k*_4,6_, and *k*_⊘,1_) are from experimental folding data reported in Refs. 6,7 under high salt conditions.

#### Analytic Expression: Ratio of the Steady-State Fluxes

The ratio of the steady-state fluxes *J_T_*/*J_B_* (see Eq. (S9)) does not depend on the energies of any of the intermediate states *ϵ_i_*. The analytic expression for this ratio of the fluxes depends only on the transition state energies 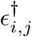:

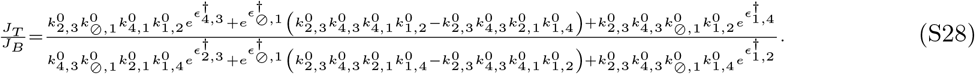

This is a consequence of Eq. (11) in the main text for the ratio of the steady-state fluxes *J_T_*/*J_B_* for the two possible pathways on the protein folding network.

### KPR Network

Table. S2 lists the kinetic parameters for the kinetic proofreading (KPR) network that describes aminoacyl(aa)-tRNA selection in the *E. coli* ribosome (see Fig. 3*A – D* in the main text). This KPR process is an important step during protein translation^8,9^, and it is performed by the ribosome with high fidelity (error rate ~ 10^−3^ − 10^−4^) ^10–14^. The biochemical network is comprised of two cycles powered by GTP hydrolysis that involve the right R (cognate)/wrong W (non-cognate) aa-tRNA molecules. This process has a product formation cycle (rates 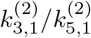) where one amino acid is added to the growing polypeptide chain to create the products P_R_/P_W_ and a proofreading cycle (rates 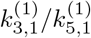) where an aa-tRNA molecule is excised from the ribosome to give R*/W*.

#### Kinetic Parameters: aa-tRNA Selection in the E. coli Ribosome

**Table S2:** Experimental Rate Constants *k_i,j_* for aa-tRNA Selection in the *E. coli* Ribosome

| System Parameters | *E. coli* Ribosome |
| --- | --- |
| $k_{1,2}$ | 40 |
| $k_{2,1}$ | 0.5 |
| $k_{2,3}$ | 25 |
| $k_{3,2}$ | $1 \times 10^{-3}$ |
| $k_{3,1}^{(1)}$ | $8.5 \times 10^{-2}$ |
| $k_{1,3}^{(1)}$ | $1 \times 10^{-3}$ |
| $k_{3,1}^{(2)}$ | 8.415 |
| $k_{1,3}^{(2)}$ | $6.9 \times 10^{-5}$ |
| $k_{1,4}$ | 27 |
| $k_{4,1}$ | 47 |
| $k_{4,5}$ | 1.2 |
| $k_{5,4}$ | $1 \times 10^{-3}$ |
| $k_{5,1}^{(1)}$ | 0.67 |
| $k_{1,5}^{(1)}$ | $2.7 \times 10^{-6}$ |
| $k_{5,1}^{(2)}$ | $3.5 \times 10^{-2}$ |
| $k_{1,5}^{(2)}$ | $9.9 \times 10^{-11}$ |

The values of the rate constants *k_i,j_* (in units of s^−1^) are from experimental data reported in Ref. 11. The chemical potential differences from Ref. 14 for the respective proofreading and the product formation cycles are fixed at their physiological values for GTP hydrolysis Δ*μ*_proof_ ~ 20 *k_B_T* and product formation Δ*μ*_p_ ~ 26 *k_B_T*. Note also that Δ*μ*_proof_ and Δ*μ*_p_ for the proofreading and the product formation cycles are equal for the right R (cognate) and wrong W (non-cognate) aa-tRNA molecules.

#### Analytic Expressions: Error Rate and Normalized Energy Dissipation

The error rate is the probability that a wrong W amino acid is added to the growing polypeptide chain. This ratio of the wrong W product formation flux *J_W_* to the right R product formation flux *J_R_* is given by:

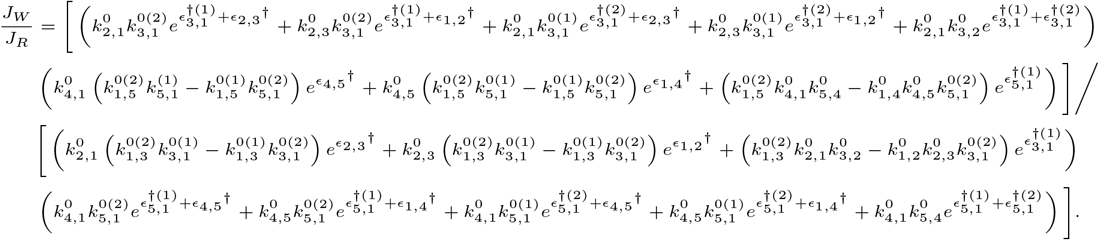

Notably, the error rate depends only on the transition state energies 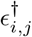, but it does not depend on the energies *ϵ_i_* of any intermediate states. As such, the error rate is invariant to energy perturbations of the intermediate states on the free energy landscape. In addition, the normalized energy dissipation *σ_N_* (see Eq. (12) in the main text) depends on the ratio of the proofreading to the product formation fluxes *J*_proof_/*J_p_*, and this property is also invariant to the energy perturbations of the intermediate states. The following analytic expression for the normalized energy dissipation depends only on the transition state energies 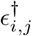:

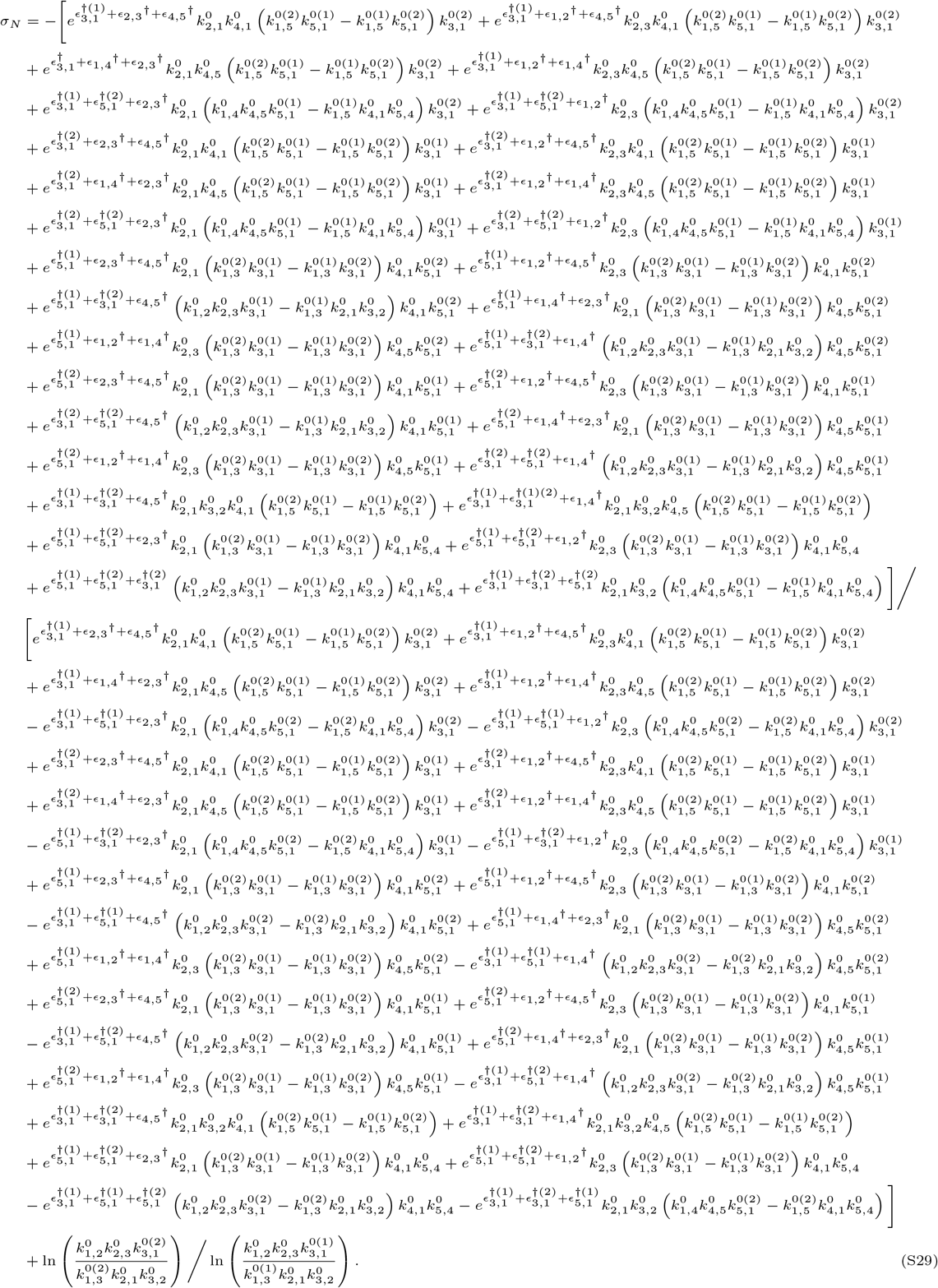

The invariance of the error rate *J_W_*/*J_R_* and the normalized energy dissipation *σ_N_* is a consequence of Eq. (11) in the main text.

### Myosin-V Network

#### Kinetic Parameters: Myosin-V Motor Protein

Table. S3 lists the kinetic parameters for the mechanochemical network that describes the motion of the myosin-V motor protein as it walks on actin cytoskeleton filaments (see Fig. 4*A – D* in the main text). This biological process is powered by energy from ATP hydrolysis, which enables the myosin-V to take one ~ 36 nm step forward an actin filament while simultaneously carrying a load that exerts an opposing force *F*_external_ on the motor protein ^4,15–19^. This kinetic model for myosin-V illustrates that the motor protein can move one step forward or backward on the actin track (via states 1-5) during a main stepping cycle or it can remain at the same position on the track during a futile cycle (via states 2, 3, 4, and 6).

**Table S3:** Rate Constants *k_i,j_* for the Myosin-V Motor Protein

| System Parameters | Myosin-V |
| --- | --- |
| $k_{1,2}$ | $1 \times 10^5$ |
| $k_{2,1}$ | 0.49 |
| $k_{2,3}$ | $4.6 \times 10^3$ |
| $k_{3,2}$ | $6.4 \times 10^4$ |
| $k_{3,4}$ | $3 \times 10^3$ |
| $k_{4,3}$ | 4.64 |
| $k_{4,5}$ | 15.2 |
| $k_{5,4}$ | 3.35 |
| $k_{5,1}$ | 302.8 |
| $k_{1,5}$ | 0.62 |
| $k_{4,6}$ | $6.9 \times 10^{-2}$ |
| $k_{6,4}$ | $1.5 \times 10^{-2}$ |
| $k_{6,2}$ | 302.8 |
| $k_{2,6}$ | $1.27 \times 10^{-6}$ |

The values of *k_i,j_* (in units of s^−1^) for this kinetic model were obtained from energetic data and kinetic parameters reported in Ref. 17. The [ATP]=[P_i_]=1 mM and [ADP]=10 *μ*M at physiological conditions, and the motor protein is carrying a load of *F*_external_ = 0.1 pN. The chemical potential differences at these physiological conditions for the futile and the main cycles respectively are Δ*μ*_futile_ ~ 25 *k_B_T* and Δ*μ*_main_ ~ 24 *k_B_T*.

#### Analytic Expressions: Ratio of the Backward- to Forward-Stepping Fluxes and Normalized Energy Dissipation

The analytic expression for the ratio of the backward-stepping flux *J_B_* to the forward-stepping flux *J_F_* of the myosin-V motor protein is given by:

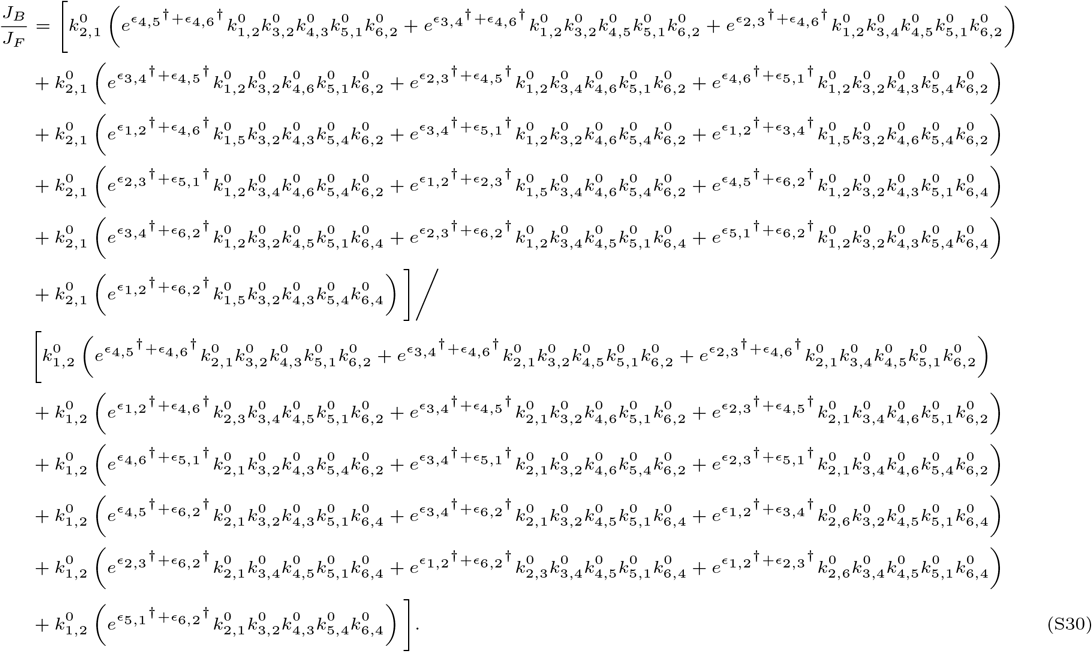

Notably, this ratio does not depend on the energies *ϵ_i_* of any intermediate states and only depends on the transition state energies 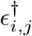. As such, this ratio of the fluxes is invariant to the energy perturbations of the intermediate states on the free energy landscape. Moreover, the normalized energy dissipation *σ_N_* (see Eq. (13) in the main text) depends on the ratio of the fluxes for the futile and the main cycles *J*_futile_/*J*_main_, and this property is also invariant to the energy perturbations of the intermediate states on the free energy landscape. The following analytic expression for the normalized energy dissipation depends only on the transition state energies 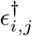:

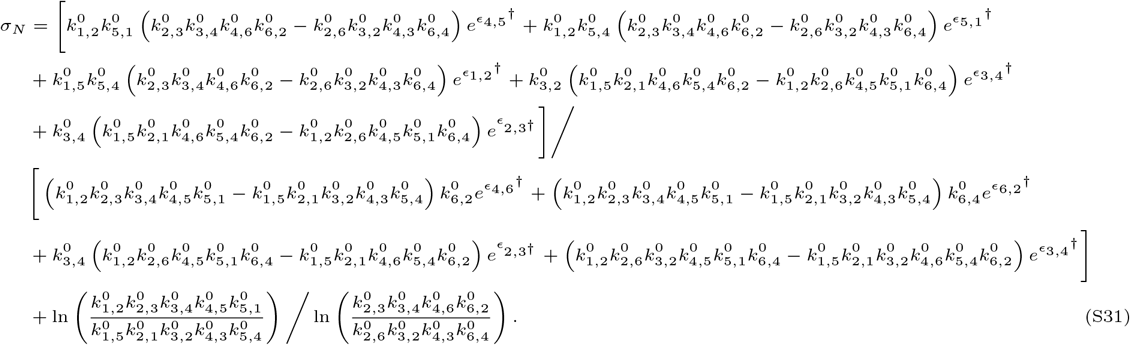

The invariance of the ratio of the backward- to forward-stepping fluxes *J_B_/J_F_* and the normalized energy dissipation *σ_N_* is a consequence Eq. (11) in the main text.

#### Energy Coupling in Protein Folding from Φ-Value Analysis

In multiple biological and chemical systems, it is frequently observed that the energy perturbations of the individual states correlate with changes in the transition-state energies. Protein folding processes provide a good example of how such energy coupling occurs and what impact it can have for interpreting the effects of mutations on the steady-state flux ratios. The extent to which the mutation perturbs the features of the underlying free-energy landscapes ofprotein folding is often determined using the so-called Φ-value analysis ^20^. Φ-value quantifies the perturbation of the transition-state free energy relative to that of the folding stability.

Conventional Φ-value analysis considers protein folding as a two-state process. In this picture, the protein can undergo folding starting from the denatured (unfolded) state *U* via a transition state (TS) or it can undergo unfolding starting from the native (folded) state *F* via the same transition state. Consequently, there are two possible definitions of the quantity Φ that can be employed in the Φ-value analysis. In the case of the folding reaction, we have

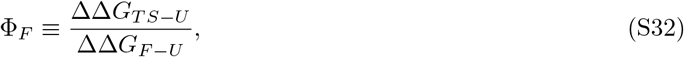

where ΔΔ*G_TS−U_* describes the change in the free energy barrier for the folding transition due to a mutation, while ΔΔ*G_F−U_* corresponds to the free energy change between the folded and unfolded protein states due to the same mutation. Similarly, in the case of the unfolding reaction (i.e., the reverse of folding), we have

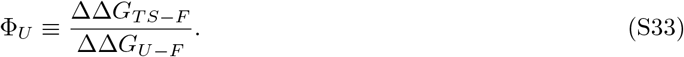

In the following discussion on multi-state protein folding processes, we have chosen to utilize Φ_*F*_ for the folding reaction.

The Φ-value analysis described above for a two-state folding process can be generalized to multi-state systems with multiple pathways and intermediate states (e.g., see the protein folding network shown in Fig. 2 of the main text). In this case, the meaning of the free energies for folded and unfolded protein states remain the same. However, the definition of the transition state free energy barrier (in units of *k_B_T*) is now effective, and it is given via the corresponding folding rates *k*_fold_:

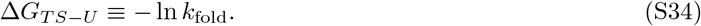

From a practical point of view, the changes in the folding kinetic rates due to the mutations are monitored and then utilized in the analysis in the same way as is done for two-state systems.

Assuming that the mutations do not modify the free energy of the denatured state *U*, perturbations that only affect the stability of the folded state *F* but not the transition-state energies, correspond to Φ_*F*_ ≈ 0. We note that these cases are not uncommon, ^21^ and our theory predicts that the ratios of steady-state fluxes (e.g., between two alternative stable folds) would be invariant to such mutations. Moreover, our invariance results can have important implications for the cases with Φ_*F*_ ≠ 0. The steady-state flux ratios (e.g., flux splitting between two different pathways) will change in the same way due to two different mutations that identically change the conformation of the transition-state. As such, these two mutations lead to the same changes in the transition-state energy barrier 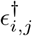. This will be the case even when Φ_*F*_ ≠ 0 and when these mutations have different effects on the stability of the individual states *ϵ_i_*. Thus, the two different mutations with Φ_*F*_ ≠ 0 can have exactly the same effect on the steady-state flux ratios even when the changes in the transition-state energies are coupled to changes in the energies of the discrete states.

These arguments demonstrate the connection of our theoretical approach with the widely utilized Φ-value analysis, and thereby, with the molecular view on of the protein folding/unfolding reactions. This connection can help researchers interpret their experimental results in light of our theory.

## References

[1] I. H. Segel, Enzyme kinetics: behavior and analysis of rapid equilibrium and steady state enzyme systems (Wiley New York:, 1975).

[2] A. Cornish-Bowden, Fundamentals of Enzyme Kinetics (John Wiley & Sons, 2013).

[3] T. Kuhlman, Z. Zhang, M. H. Saier, and T. Hwa, “Combinatorial transcriptional control of the lactose operon of escherichia coli,” Proc. Natl. Acad. Sci. USA 104, 6043–6048 (2007).

[4] H. D. Kim and E. K. O’shea, “A quantitative model of transcription factor–activated gene expression,” Nat. Struct. Mol. Biol. 15, 1192 (2008).

[5] J. Narula and O. Igoshin, “Thermodynamic models of combinatorial gene regulation by distant enhancers,” IET Syst. Biol. 4, 393–408 (2010).

[6] J. Estrada, F. Wong, A. DePace, and J. Gunawardena, “Information integration and energy expenditure in gene regulation,” Cell 166, 234–244 (2016).

[7] K. L. Pierce, R. T. Premont, and R. J. Lefkowitz, “Seven-transmembrane receptors,” Nat. Rev. Mol. Cell Biol. 3, 639 (2002).

[8] J.-P. Changeux and S. J. Edelstein, “Allosteric mechanisms of signal transduction,” Science 308, 1424–1428 (2005).

[9] D. Colquhoun, “Agonist-activated ion channels,” Br. J. Pharmacol. 147, S17–S26 (2006).

[10] S. Prabakaran, G. Lippens, H. Steen, and J. Gunawardena, “Post-translational modification: nature’s escape from genetic imprisonment and the basis for dynamic information encoding,” Wiley Interdiscip. Rev. Syst. Biol. Med. 4, 565–583 (2012).

[11] J. Gunawardena, “Time-scale separation–michaelis and menten’s old idea, still bearing fruit,” FEBS J. 281, 473–488 (2014).

[12] J. Gunawardena, “A linear framework for time-scale separation in nonlinear biochemical systems,” PloS One 7, e36321 (2012).

[13] D. E. Makarov, Single Molecule Science: Physical Principles and Models (CRC Press, 2015).

[14] K. J. Laidler, Chemical Kinetics (1987).

[15] T. L. Hill, Free Energy Transduction and Biochemical Cycle Kinetics (Courier Corporation, 2005).

[16] G. K. Ackers, A. D. Johnson, and M. A. Shea, “Quantitative model for gene regulation by lambda phage repressor,” Proc. Natl. Acad. Sci. USA 79, 1129–1133 (1982).

[17] L. Bintu, N. E. Buchler, H. G. Garcia, U. Gerland, T. Hwa, J. Kondev, and R. Phillips, “Transcriptional regulation by the numbers: models,” Curr. Opin. Genet. Dev. 15, 116–124 (2005).

[18] P. Sartori and S. Pigolotti, “Kinetic versus energetic discrimination in biological copying,” Phys. Rev. Lett. 110, 188101 (2013).

[19] J. J. Hopfield, “Kinetic proofreading: A new mechanism for reducing errors in biosynthetic processes requiring high specificity,” Proc. Natl. Acad. Sci. 71, 4135–4139 (1974).

[20] J. Howard, “Molecular motors: structural adaptations to cellular functions,” Nature 389, 561–567 (1997).

[21] A. Murugan, D. A. Huse, and S. Leibler, “Discriminatory proofreading regimes in non-equilibrium systems,” Phys. Rev. X 4, 021016 (2014).

[22] A. B. Kolomeisky, Motor Proteins and Molecular Motors (CRC Press, Taylor and Francis Group: Boca Raton, FL, 2015).

[23] J. Ninio, “Kinetic amplification of enzyme discrimination,” Biochimie 57, 587–595 (1975).

[24] K. I. Skau, R. B. Hoyle, and M. S. Turner, “A kinetic model describing the processivity of myosin-V,” Biophys. J. 91, 2475–2489 (2006).

[25] K. Banerjee, A. B. Kolomeisky, and O. A. Igoshin, “Accuracy of substrate selection by enzymes is controlled by kinetic discrimination,” J. Phys. Chem. Lett. 8, 1552–1556 (2017).

[26] S. W. Englander, L. Mayne, and M. M. Krishna, “Protein folding and misfolding: mechanism and principles,” Q. Rev. Biophys. 40, 1–41 (2007).

[27] M. M. Krishna and S. W. Englander, “A unified mechanism for protein folding: predetermined pathways with optional errors,” Protein Sci. 16, 449–464 (2007).

[28] H. S. Zaher and R. Green, “Fidelity at the molecular level: lessons from protein synthesis,” Cell 136, 746–762 (2009).

[29] N. M. Reynolds, B. A. Lazazzera, and M. Ibba, “Cellular mechanisms that control mistranslation,” Nat. Rev. Microbiol. 8, 849 (2010).

[30] K. Banerjee, A. B. Kolomeisky, and O. A. Igoshin, “Elucidating interplay of speed and accuracy in biological error correction,” Proc. Natl. Acad. Sci. USA 114, 5183–5188 (2017).

[31] J. D. Mallory, A. B. Kolomeisky, and O. A. Igoshin, “Trade-offs between error, speed, noise, and energy dissipation in biological processes with proofreading,” J. Phys. Chem. B 123, 4718–4725 (2019).

[32] T. H. Lowry and K. S. Richardson, Mechanism and Theory in Organic Chemistry (Harper & Row, 1987).

[33] O. Bieri, G. Wildegger, A. Bachmann, C. Wagner, and T. Kiefhaber, “A salt-induced kinetic intermediate is on a new parallel pathway of lysozyme folding,” Biochemistry 38, 12460–12470 (1999).

[34] H. S. Zaher and R. Green, “Hyperaccurate and errorprone ribosomes exploit distinct mechanisms during tRNA selection,” Mol. Cell 39, 110–120 (2010).

[35] C. M. Dobson, “Protein folding and misfolding,” Nature 426, 884 (2003).

[36] A. Berezhkovskii, G. Hummer, and A. Szabo, “Reactive flux and folding pathways in network models of coarsegrained protein dynamics,” J. Chem. Phys. 130, 05B614 (2009).

[37] S. B. Prusiner, “Prions,” Proceedings of the National Academy of Sciences 95, 13363–13383 (1998).

## References

[1] J. Gunawardena, “A linear framework for time-scale separation in nonlinear biochemical systems,” PloS One 7, e36321 (2012).

[2] T. L. Hill, Free Energy Transduction and Biochemical Cycle Kinetics (Courier Corporation, 2005).

[3] M. E. Tuckerman, Statistical Mechanics: Theory and Molecular Simulation (Oxford University Press, 2010).

[4] A. B. Kolomeisky, Motor Proteins and Molecular Motors (CRC Press, Taylor and Francis Group: Boca Raton, FL, 2015).

[5] S. W. Englander, L. Mayne, and M. M. Krishna, “Protein folding and misfolding: mechanism and principles,” Q. Rev. Biophys. 40, 1–41 (2007).

[6] M. M. Krishna and S. W. Englander, “A unified mechanism for protein folding: Predetermined pathways with optional errors,” Protein Sci. 16, 449–464 (2007).

[7] O. Bieri, G. Wildegger, A. Bachmann, C. Wagner, and T. Kiefhaber, “A salt-induced kinetic intermediate is on a new parallel pathway of lysozyme folding,” Biochemistry 38, 12460–12470 (1999).

[8] K. B. Gromadski and M. V. Rodnina, “Kinetic determinants of high-fidelity tRNA discrimination on the ribosome,” Mol. Cell 13, 191 (2004).

[9] N. M. Reynolds, B. A. Lazazzera, and M. Ibba, “Cellular mechanisms that control mistranslation,” Nat. Rev. Microbiol. 8, 849 (2010).

[10] H. S. Zaher and R. Green, “Fidelity at the molecular level: lessons from protein synthesis,” Cell 136, 746–762 (2009).

[11] H. S. Zaher and R. Green, “Hyperaccurate and error-prone ribosomes exploit distinct mechanisms during tRNA selection,” Mol. Cell 39, 110–120 (2010).

[12] K. Banerjee, A. B. Kolomeisky, and O. A. Igoshin, “Elucidating interplay of speed and accuracy in biological error correction,” Proc. Natl. Acad. Sci. USA 114, 5183–5188 (2017).

[13] K. Banerjee, A. B. Kolomeisky, and O. A. Igoshin, “Accuracy of substrate selection by enzymes is controlled by kinetic discrimination,” J. Phys. Chem. Lett. 8, 1552–1556 (2017).

[14] J. D. Mallory, A. B. Kolomeisky, and O. A. Igoshin, “Trade-offs between error, speed, noise, and energy dissipation in biological processes with proofreading,” J. Phys. Chem. B 123, 4718–4725 (2019).

[15] M. Enrique, A. L. Wells, S. S. Rosenfeld, E. M. Ostap, and H. L. Sweeney, “The kinetic mechanism of myosin V,” Proc. Natl. Acad. Sci. USA 96, 13726–13731 (1999).

[16] A. B. Kolomeisky and M. E. Fisher, “A simple kinetic model describes the processivity of myosin-V,” Biophys. J. 84, 1642–1650 (2003).

[17] K. I. Skau, R. B. Hoyle, and M. S. Turner, “A kinetic model describing the processivity of myosin-V,” Biophys. J. 91, 2475–2489 (2006).

[18] A. D. Mehta, R. S. Rock, M. Rief, J. A. Spudich, M. S. Mooseker, and R. E. Cheney, “Myosin-V is a processive actin-based motor,” Nature 400, 590 (1999).

[19] A. Mehta, “Myosin learns to walk,” J. Cell Sci. 114, 1981–1998 (2001).

[20] A. R. Fersht and S. Sato, “Φ-value analysis and the nature of protein-folding transition states,” Proceedings of the National Academy of Sciences USA 101, 7976–7981 (2004).

[21] W. Paslawski, O. K. Lillelund, J. V. Kristensen, N. P. Schafer, R. P. Baker, S. Urban, and D. E. Otzen, “Cooperative folding of a polytopic *α*-helical membrane protein involves a compact n-terminal nucleus and nonnative loops,” Proceedings of the National Academy of Sciences 112, 7978–7983 (2015)

